# WASP facilitates tumor mechanosensitivity in T lymphocytes

**DOI:** 10.1101/2023.10.02.560434

**Authors:** Srishti Mandal, Mariane Melo, Pavlo Gordiichuk, Sayanti Acharya, Yeh-Chuin Poh, Na Li, Aereas Aung, Eric L. Dane, Darrell J. Irvine, Sudha Kumari

**Affiliations:** Indian Institute of Science, Bengaluru, India; Koch Institute of Integrative Cancer Research, MIT, Cambridge, USA; Department of Biological Engineering, MIT, Cambridge, USA; Howard Hughes Medical Institute, Ashburn, Virginia, USA

**Keywords:** Mechanosensing, Cytotoxic T cell activation, Tumor cytotoxicity, Actin cytoskeleton, Wiskott-Aldrich Syndrome Protein

## Abstract

Cytotoxic T lymphocytes (CTLs) carry out immunosurveillance by scanning target cells of diverse physical properties for the presence of antigens. While the recognition of cognate antigen by the T cell receptor is the primary signal for CTL activation, it has become increasingly clear that the mechanical stiffness of target cells plays an important role in antigen-triggered T cell responses. However, the molecular machinery within CTLs that transduces the mechanical information of tumor cells remains unclear. We find that CTL’s mechanosensitive ability requires the activity of the actin-organizing protein Wiskott-Aldrich Syndrome Protein (WASP). WASP activation is modulated by the mechanical properties of antigen-presenting contexts across a wide range of target cell stiffnesses and activated WASP then mediates mechanosensitive activation of early TCR signaling markers in the CTL. Our results provide a molecular link between antigen mechanosensing and CTL immune response and suggest that CTL-intrinsic cytoskeletal organizing principles enable the processing of mechanical information from diverse target cells.

## Introduction

T cells search for antigens by physically scanning the surface of target cells. If a cognate antigen peptide displayed on MHC molecules (pMHC) of target cells is detected by the T cell Receptor (TCR), a dynamic cell-cell junction termed an ‘immunological synapse’ forms between the target cell and the T cell^1^. In the case of cytotoxic T lymphocytes (CTLs), the synaptic interface serves to provide both-the recognition and initial activation signal to the TCR as well as the physical site where CTLs deliver cytotoxic molecules to kill infected or malignant target cells. It has been shown that the mechanical forces at the CTL synapse influence both the early activation of T cells as well as their subsequent cytotoxic function^2–15^. Despite mounting evidence that the mechanical context of antigen presentation dictates T cell activation and subsequent immune response^7,13^, the mechanisms by which mechanical forces at the synapse are sensed and transduced within the CTLs are not completely clear. Recent studies have shown that TCR can participate in the mechanosensitive activation of T cells by utilizing forces for improving the lifetime of the bond with the MHC-antigen ^8,12,16,17^ and by involving as yet uncharacterized downstream mechanisms at the cell surface resulting in mechanosensitivity^18–20^. Still, the molecular pathways downstream of TCR that consolidate and transduce the mechanical information to the cell interior remain unclear.

Experiments using pharmacological perturbation of actin nucleation factors suggest that the mechanosensing activity of the antigen receptor is supported by the cortical actin cytoskeleton within the lymphocytes ^5,9,11^,^21–28^. However, molecular mechanisms within the lymphocytes that connect antigen-presenting surface mechanics to the underlying cytoskeleton remain unknown. In addition, whether the target cell cytoskeleton is also involved in lymphocyte mechanosensing is not known. This is a crucial knowledge gap, since cortical cytoskeletal processes would be the primary modulators of target cell stiffness^29^ and would thereby regulate lymphocyte immune response^2,7,14,22,30–34^.

Using primary lymphocytes activated by APC-mimetic substrates or live tumor cells, we find that mechanosensitive recognition of target cells by CTLs is mediated by WASP. As a crucial regulator of the actin polymerization program at the synapse^35–41^, WASP leverages target cell stiffness to potentiate CTL responses such that antigen-triggered CTL activation, and eventually cytotoxicity and anti-tumor activity are all modulated by WASP as the target cell stiffness changes. These results reveal a T cell-intrinsic cytoskeletal program that underlies mechanical crosstalk between the CTL and tumor cells and allows successful target recognition and destruction. These results raise the possibility that modulation of both CTL as well as tumor cortical stiffness via specific actin nucleation programs could be a candidate mechanism for tumor cell evasion from optimal recognition and cytotoxicity by killer T cells. Our study also sheds light on immune cytoskeletal mechanisms that are dysregulated in cancer and could contribute to the high incidence of cancer seen in WAS immunodeficient individuals.

## Results and discussion

For effector function, CTLs first scan target cells of diverse mechanical properties for the presence of antigens. The CTL activation process relies significantly on the mechanical stiffness of target cells during the antigen recognition process. Actin cytoskeleton is known to support mechanosensitive T cell activation^10^, and since Wiskott –Aldrich Syndrome Protein (WASP) is a major actin regulator in T cells, we wondered if WASP enables T cell antigen mechanosensing.

To establish a cellular system of differential antigen mechanosensing, we first modulated the target tumor cell’s cortical stiffness using non-reversible pharmacological inhibition of the actin cytoskeleton. The B16F10 (B16) melanoma tumor cells were acutely treated with the irreversible F-actin inhibitor mycalolide B (Myca.) for 10 min followed by washing to remove unbound inhibitor^42^, and atomic force microscopy (AFM)-based stiffness measurement **(Fig. 1A, B)**. Myca. treatment of B16 cells induced a ~50% reduction in their elastic modulus **(Fig. 1C)**, whereas surface expression of H-2D^b^-the MHCI molecule crucially required for antigen presentation and thereby early T cell activation in this system-remained indistinguishable from the DMSO-treated control group **(Fig. S1A, B)**. To assess tumor antigen-triggered TCR activation, we employed CTLs derived either from Wild Type (WT) or from WASP-/- TCR-transgenic pmel-1 mice (the pmel-1 TCR is specific for a peptide from gp100 in the context of H-2D^b^ on B16 cells)^43^. The WT or WASP-deficient pmel-1 CTLs were incubated with Myca.-pretreated or control B16 cells for 5 min to allow immunological synapse formation, assembly of filamentous actin (F-actin) at the synapse, and TCR activation. The synaptic interface between WT and WASP-/- CTLs and B16 targets showed an accumulation of filamentous actin (“Actin”), indicating that global F-actin assembly proceeds normally in CTLs interacting with control or Myca.-treated “softened” B16 cells (Fig.1D, upper panels). However, we observed organizational differences in the F-actin at the synapse. The WT CTL-control B16 conjugates showed actin foci across the synaptic interface, which were missing in WASP^-/-^ CTL-control B16 synapses. A similar lack of actin foci was also observed in WT CTL -Myca. B16 synapses **(Fig. 1D, lower panels, graph)**. The WASP-dependent foci-like structures observed in WT CTL -control B16 synapses were highly dynamic **(Movie1)** and were reminiscent of WASP-dependent actin foci described previously in CD4^+^ T cells, which contribute to T cell activation^44,45^. Thus, we next examined the effect of B16 softening on early TCR signaling. We assessed the phosphorylation of the TCR CD3ζ chain in one of its ITAM motifs (Tyr 149)^46,47^ and of the adaptor protein Zap70 at Tyr 319^48^ in the CTL-tumor cell synaptic interface. These early TCR signaling markers were reduced in both WASP-/- CTL-control B16 as well as WT CTL – Myca. B16 synapses to a similar extent (**Fig. 1E**). The similarities in actin organization and early TCR signaling defects when either CTLs lack WASP, or B16 targets are soft suggested that sensing of target cell stiffness might be linked to WASP activity. Consistently, WT CTL-Myca-B16 synapses showed significantly lower WASP activation, as marked by levels of Y290-phosphorylated WASP (pWASP) at the interface **(Fig. 1F)**. The reduced WASP activation and phosphorylation of early TCR signaling markers was further associated with poor cytotoxicity in the *in vitro* cytolysis assay **(Fig. 1G)**. While WT CTLs incubated with control B16 cells exhibited potent cytotoxicity, both WT CTLs incubated with Myca.-treated B16 cells and WASP-/- CTLs incubated with control B16 cells showed severely impaired cytotoxic activity **(Fig. 1G)**.

**Figure 1.**
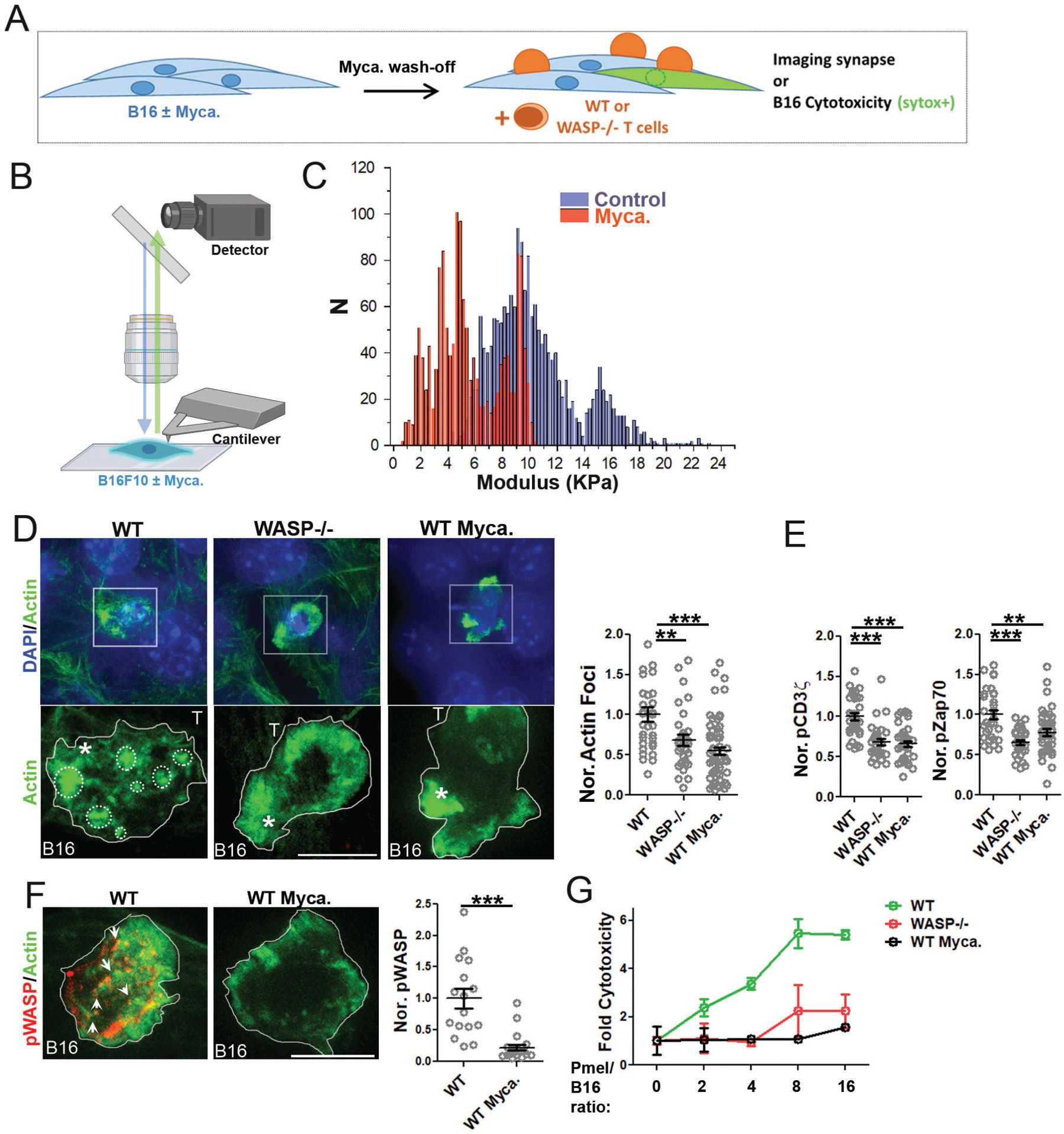
Acute Myca. treatment induces tumor cell softness, reduction in early signaling markers, and a loss of actin foci and WASP activation in tumor-CTL synapses. (A) Schematic of the pre-treatment of B16 cells with DMSO or 1µM Mycalolide A (Myca) in B16-CTL co-culture assays. (B) AFM setup for B16 stiffness measurements. (C) comparison of DMSO (control) or Myca.-treated B16 stiffness. Each bar in the plot represents the value obtained from a single cell, the p-value of the distributions is <0 .00001, as determined using Mann-Whitney two-tailed test. (D-G) Effects of Myca treatment on events at the B16-CTL synapses. Cells were fixed after 5 min for confocal imaging (D-F) or cultured for 4 h followed by assessment of target cell killing by Sytox staining (G). Images in (D) and (F) show the maximum intensity projected *<u>en-face</u>* view of F-actin in the CTL/target interfaces. The actin foci are demarcated in white dotted circles (D, left panel), or arrows (F, left panel). The graphs show quantification of F-actin foci (D), phospho-CD3ζ and Zap70 (E), and phospho-WASP (F). Each point in the scatter plot represents the values obtained from individual cells; the bars show Mean± SEM. These experiments were repeated at least thrice with similar results. (G) *In vitro* killing of B16F10 target cells as a function of Effector: Target (E: T) ratio. Each data point represents values obtained from four independent samples as Mean ± SEM. This experiment was repeated twice with similar results. Scale bar in the images, 5µm.

An explanation of the result that Myca.-treated B16 cells instigate poor CTL activation and cytotoxic response could be that Myca. leaches outside of tumor cells into co-culture media and inhibits the CTL cytoskeleton. To test if indeed Myca. leaching from antigen-bearing cells could account for poor T cell activation, we utilized the bone marrow-derived dendritic cell (BMDC) and OTII CD4+ T cell co-culture system. This system allows testing of the short-term and long-term effects of antigen presenting cell-induced T cell activation that is not possible in the B16-CTL system. In this assay, pre-treatment of BMDCs with Myca. Followed by a washout led to poor activation of OTII cells characterized by impaired modulation of surface markers CD69 and CD62L, effector cytokine production, and activation-induced proliferation (**Fig. S2A-F**). All of these defects were significantly restored if mock-treated BMDCs were also present in the wells. This indicates that the leaching of Myca. from treated cells does not cause inhibition of bystander cells and is unlikely to be the reason for the poor CTL activation by Myca-treated B16 cells. Together, these results indicate that Myca-induced changes in the mechanical stiffness of tumor cells impact CTL activation, and WASP could provide a molecular connection between them.

There could be two possible explanations for the defective cytotoxic function of WASP-/- CTLs. It could be due to a loss of WASP-dependent actin polymerization, or because of a loss of WASP-based protein-protein interactions at the synapse that would have allowed TCR activation to proceed. To tease apart these two possibilities, we expressed a mutant form of WASP lacking only 8 amino acids in its c-terminal domain (WASPΔC). WASPΔC cannot activate the Arp2/3 complex and thereby initiate actin polymerization but should perform the other protein scaffolding roles of WASP^44^. Expression of WASPΔC led to reduced actin enrichment at the synaptic interface with a simultaneous reduction in TCR activation and CTL cytotoxic activity *in vitro*. **(Fig. 2A-B)** The similarity in WASP-/- and WASPΔC CTL phenotypes indicated that WASP’s nucleation-promoting factor (NPF) activity is essential for its support of TCR activation and CTL cytotoxicity. To further test whether the mechanical stresses are compromised in WASP-/- CTL synapses, we utilized the lymphocyte form of P130Cas (CasL), phosphorylated at Tyr-249, as endogenous mechanotransduction sensor^30^. P130 Cas as well as CasL localize to filamentous actin and show higher phosphorylation at Tyr-249 in response to elevated local cytoskeletal stresses^30,45,49^. Indeed, phospho-Tyr-249-CasL (pCasL) was reduced in the synapses formed between B16 and WASP-/- CTL and could be significantly rescued by the expression of full-length WASP protein. These results indicate that cortical cytoskeletal stresses are compromised in WASP-/- CTLs. Reconstitution of WASP also significantly rescued both actin organization into foci as well as cytotoxicity in WASP^-/-^ CTLs **(Fig 2C-D)**.

**Figure 2.**
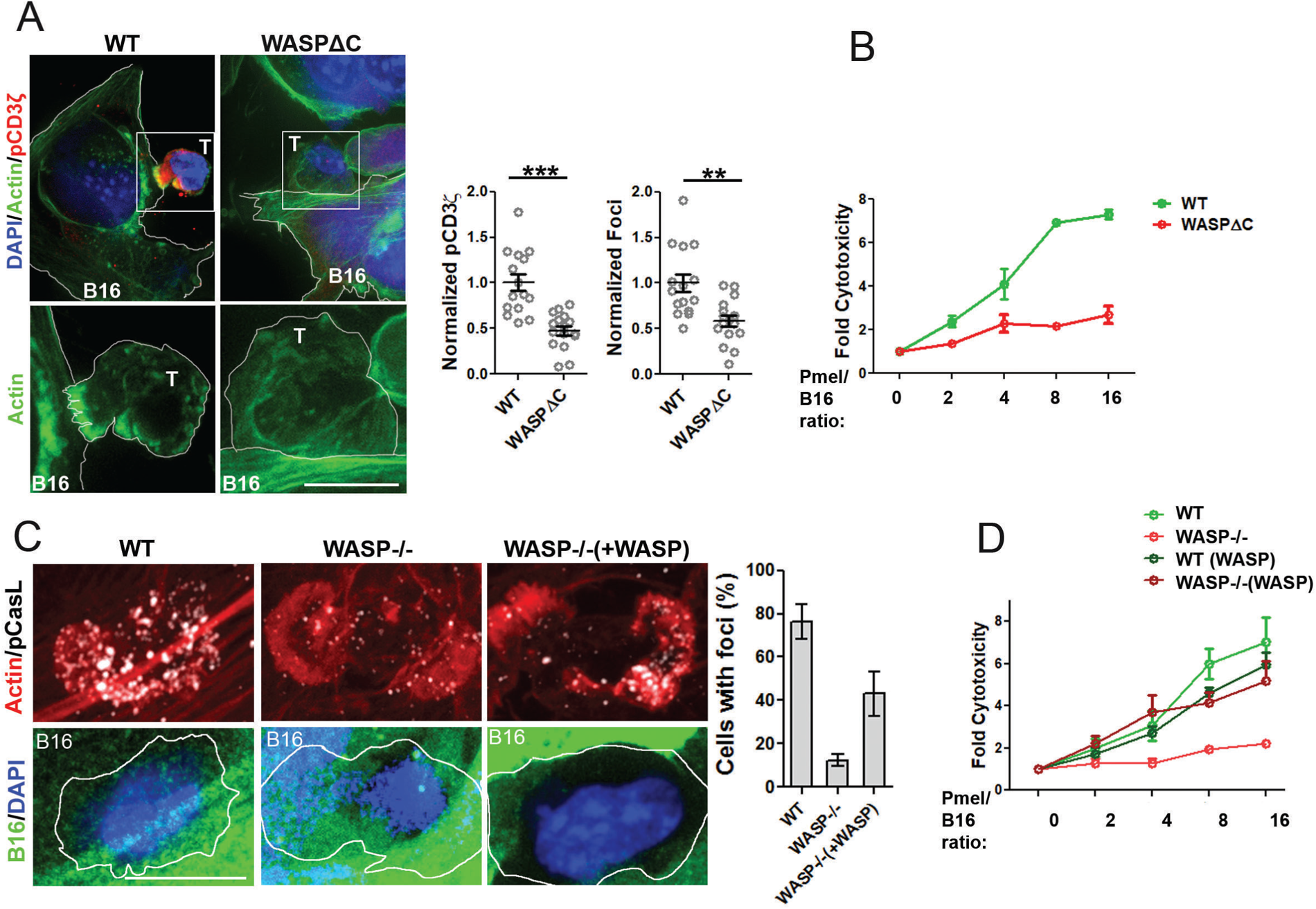
(A-B) NPF activity of WASP is required for TCR activation at B16-CTL synapse and CTL cytotoxicity. The actin foci, associated mechanical stress, and cytotoxicity can be restored by overexpression of WASP in WASP-/- CTLs. (A) Confocal images showing a *<u>s</u>*<u>ide view</u> of F-actin in the B16-pmel-1 cell synapses. Pmel-1 cells transfected with either GFP-WASP (WT) or GFP-WASPΔC (WASPΔC) were allowed to form synapses with B16 cells for 5min. The graphs show F-actin foci and phospho-CD3ζ intensity at B16-pmel-1 synapses. Each point in the scatter plot represents the values obtained from individual cells; the bars show Mean± SEM. This experiment was repeated thrice with similar results. p-values were obtained using Mann-Whitney two-tailed test; p***<0.0001, and **<0.005. (B) Cytotoxic activity of WT or WASPΔC expressing pmels. The data points in the graph represent Mean ± SEM from four replicates. (C) Images showing actin foci and pCasL levels in synapses formed between B16 cells and WT or WASP-/- pmel-1 CTLs, either expressing empty control vector (‘WT or WASP-/-’) or WASP construct. The graph shows the percentage of cells with discernable actin foci in the synapse. (D) Cytotoxic activity of WASP overexpressing pmels compared to the WT or WASP-/- pmels. The experiment was conducted as described in (B). Scale bar in the images, 5µm.

Another possible explanation for the poor cytotoxicity of WASP-/- CTLs could be the impaired release of cytotoxic molecules at the synapse in the absence of WASP. However, this turned out not to be the case as TCR-activated secretory protease release **(Fig. S3A)** or surface marker LAMP1 cycling **(Fig. S3B)** were not reduced in WASP-/- CTLs, consistent with previous studies^35^.

Treatment with Myca. led to tumor cell softening and poor TCR-induced CTL activation. However, as Myca. treatment appears to cause a bimodal shift in B16 stiffness, the cellular assay is limited in its range of mechanical perturbations. To characterize the WASP-dependent mechanosensitivity of CTLs across a broader range of stiffness, we activated CTLs on polyacrylamide (PA) gels of varying stiffnesses. The gels were surface-functionalized with a constant density of anti-CD3 antibody and ICAM-1, and Young’s moduli of the gels varied from 2.5 to 50KPa to span the stiffness range of target cells encountered by CTLs in physiological situations^14,50^. The gel substrates contained fluorescent beads embedded within the matrix at a density of ~25 beads/100 µm^2^ (**Figs. S4A-B**). WT or WASP^-/-^ CTLs were seeded onto these substrates, and forces exerted by the cells on the substrate were calculated by traction force microscopy (TFM)^51^ by recording fluorescent bead displacements within the gels during the first 5 minutes of CTL-PA gel encounter (**Fig. 3A**). After seeding, WT and WASP^-/-^ CTLs showed similar adhesion, although in both cases more cells adhered to substrates of higher stiffness^14^ (**Fig. S4C**). Adherent WT CTLs displayed a substrate stiffness-dependent increase in traction forces, which was dampened in WASP-deficient CTLs, which generated 2-3-fold lower traction forces (**Fig. 3B**) **(Movie 2-3)**. Immunostaining to visualize the synapses formed by CTLs engaged with these surfaces revealed higher Zap70 kinase phosphorylation^48^, adaptor protein Talin recruitment, and overall filamentous actin (F-actin) accumulation at the T cell-substrate interface as the substrate stiffness increased; this mechanoresponse was also substantially reduced in WASP-/- CTLs (**Fig. 3C, D)**. Like the case of CTL synapses formed with stiffer B16 cells, higher stiffness PA gel substrates induced better WASP activation and actin foci in the interface (**Fig. 3C-arrows in the image, Fig. 3E,F**). These data indicate that during the early phases of T cell activation, CTLs sense the mechanical rigidity of the substrate and transduce this information for further mechanical force generation, TCR activation and early signaling events in a WASP-dependent fashion. To better establish the relationship between WASP activity and TCR phosphorylation, we examined CTL – substrate synapses at higher magnification. While the WT CTLs displayed abundant actin foci on PA substrates that were significantly co-localized with pCD3ζ, in CTLs lacking WASP, both clustering as well as the levels of pCD3ζ were significantly diminished. As a control, the localization of integrin adaptor protein Talin with actin foci was significantly lower when compared to the localization of pCD3ζ with actin foci **(Fig. S5A)**^52^.

**Figure 3.**
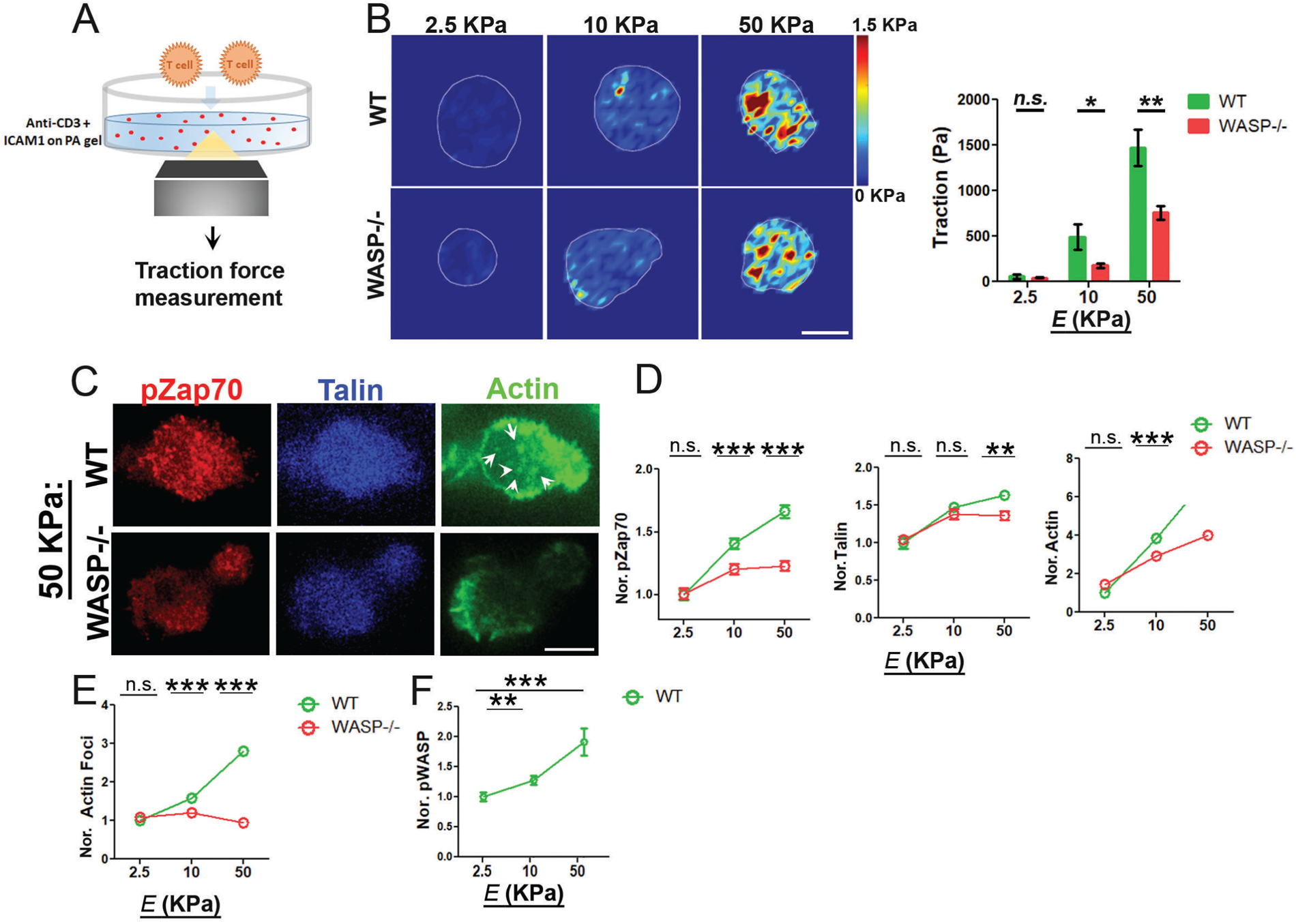
CTLs show an increased generation of mechanical forces and activation on stiffer substrates in a WASP-dependent manner. (A) Experimental setup of the traction force measurements during CTL activation on gels covalently functionalized with anti-CD3 and ICAM-1. (B) Shows representative images of traction force maps in CTL-substrate interfaces, and traction force values as a function of substrate stiffness (the graph on the right) p values, ns>0.5, *=0.03 **=0.007. (C-F) The levels of pZap70, Talin, F-actin, F-actin foci, and pWASP intensities at the CTL-substrate interfaces. In graphs (D-F) the points represent mean ± SEM values. p-values were obtained using Mann-Whitney two-tailed test by comparison of values from WT and WASP-/- cells for each stiffness; p**=0.007, and *=0.03. These experiments were repeated at least twice with similar results.

To investigate if WASP-dependent mechanosensing in early TCR signaling events eventually impacts distal events of CTL activation as well, we activated CTLs using plate-bound anti-CD3/CD28-a routine method of T cell activation using high rigidity substrates. As a control, we examined responses of CTLs to pharmacological activation by PMA/ionomycin (P/I) that can trigger CTL activation independent of TCR engagement. Following stimulation, although both WT and WASP^-/-^ cells showed comparable surface levels of the integrin LFA-1 that is crucial for the interaction of CTLs with target cells^35^ (**Fig. 4A**), WASP-deficient CTLs showed an impaired activation-induced increase in activation marker CD69 (**Fig. 4B**), secretion of the effector cytokines IL-2 **(Fig. 4C)** and IFN-γ (**Fig. 4D**), and proliferation (**Fig. 4E**). These effects were specific to anti-CD3/28-triggered activation, as pharmacological activation by P/I^53^ led to a comparable activation of distal events in WT and WASP^-/-^ CTLs (**Fig. 4B-E**). These results indicate that at high rigidity, WASP^-/-^CTLs have defective distal TCR activation responses arising from defects in early TCR signaling events since they can be activated normally by pharmacological reagents that bypass early TCR signaling. To finally establish if the role of WASP in early and late TCR signaling events regulates CTLs’ anti-tumor effector function *in vivo*, we deployed a subcutaneously implanted tumor growth assay **(Fig. 4F)**. B16 cells were injected in WT or WASP deficient mice and tumor growth was monitored. WASP-/- mice developed tumors at a much faster rate than in WT mice **(Fig. 4G)**. WASP is expressed by all immune cells, and many immune cell types in principle could contribute to higher tumor development in WASP-/- animals. To ascertain that the increased tumor growth rate is contributed by defects in CTL activity in WASP-/- animals, we then utilized adoptive therapy models of CTL function. WT C57BL/6 mice were implanted with either B16 or chicken ovalbumin peptide expressing B16 (B16-ova) tumor cells. The mice were then injected with WT or WASP-/- CTLs derived from either pmel-1 or OT1 transgenic mice for therapy **(Fig. 4H)**. In both tumor models, WASP-/- CTLs proved to be significantly impaired in their ability to limit tumor growth when compared to their WT CTL counterparts **(Fig. 4I-J)**. Collectively, these results show that WASP is also required for optimal CTL anti-tumor effector function *in vitro* as well as *in vivo*.

**Figure 4.**
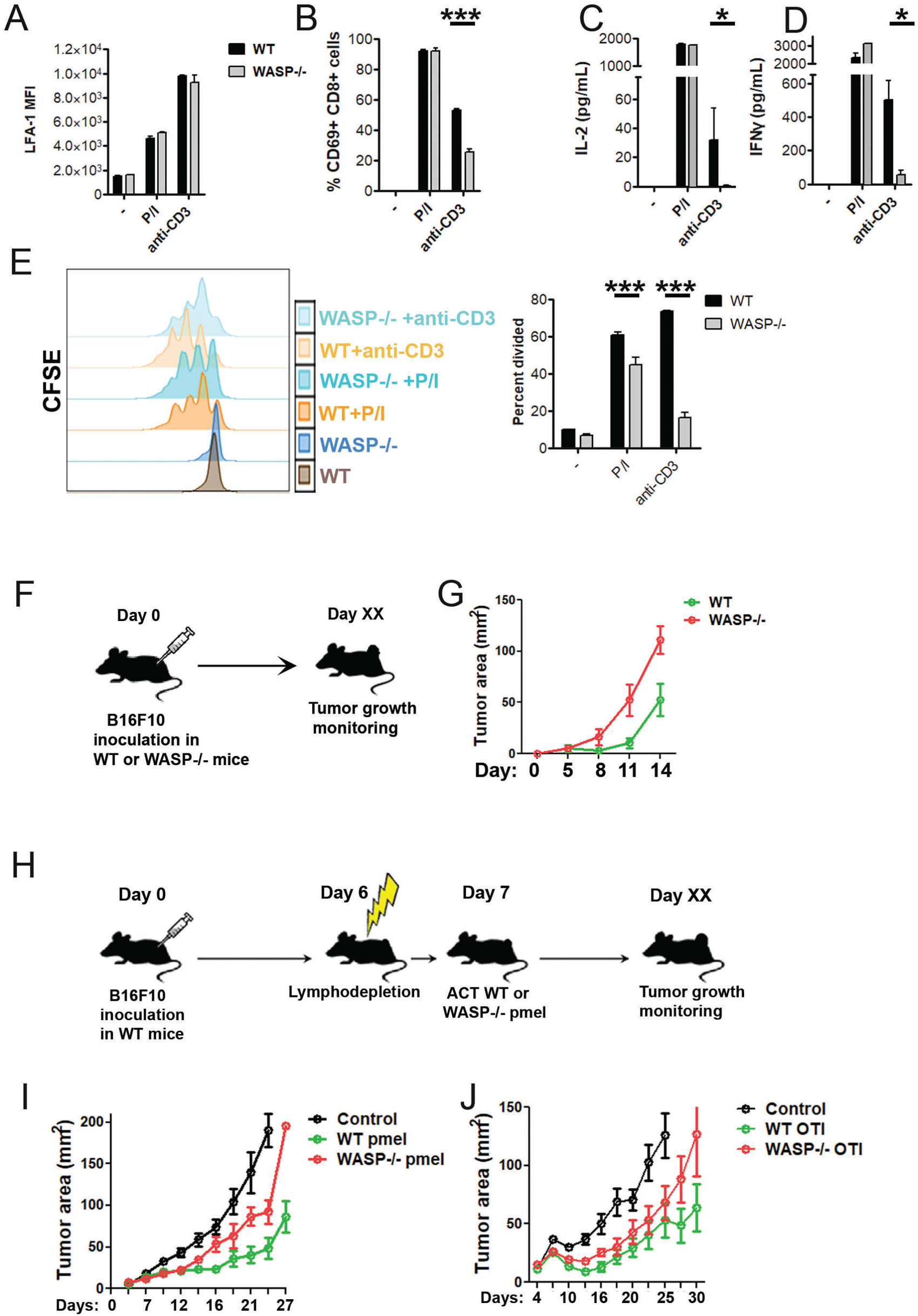
WASP-dependent CD8+ T cell activation, and *in-vivo* anti-tumor efficacy of WASP-deficient CTLs. Surface expression of LFA-1 (A), CD69 (B), or cytokine production (C-D) was assessed after activation of CD8+T cells with indicated reagents for 24h. (E) The activation-induced proliferation of CD8+T cells was determined using a CFSE dilution assay. All these experiments were repeated at least three times with similar results. The bar graphs represent mean ± SEM. *p-value* *<0.05; ***<0.005, as determined by paired two-way t-test. (F, G) C57BL/6 mice lacking WASP in their immune cells show significantly higher growth of tumors. WT or WASP-/- animals were subcutaneously injected with 0.5 million B16 cells. Tumor growth was measured using the calipers method at the indicated time points. Each data point represents data from at least six animals, and the experiment was repeated four times. (H) Adoptive transfer of WT or WASP-/- pmel-1 CTLs (left graph) or OTI CTLs in animals bearing B16 (left panels) or B16F10-ova (right panels) tumors respectively. Tumor growth was measured and plotted as described in (I-J). This experiment was repeated thrice with similar results.

## Discussion

Cytotoxic T cells utilize forces to potentiate their activation program and cytotoxic function; however, the molecular machinery within CTLs that enables the transduction of the mechanical information of the target cells is not known. Here we show, for the first time to our knowledge, that actin organizing protein WASP in CTL is required for sensing target cell rigidity. In addition, our data provide evidence that the CTL mechanosensitive response proceeds via crosstalk between cytoskeletal forces of the CTLs and the actin cytoskeleton of target tumor cells.

Our study shows that CTLs enhance their activation and cytotoxicity potential in response to an increase in target cell rigidities by increasing WASP activation. The structural basis of the mechanosensitive WASP activation and precisely how the WASP-generated cytoskeleton could promote mechanotransduction is not yet clear. The mechanosensing property could emerge from force-dependent alterations in WASP conformation leading to its activation, or a load-adapting function of actin organization (“Foci”) generated by WASP. Such an effect of forces on the density of the branched actin network has been reported previously^54^. In addition, actin foci are dynamic structures that undergo formation, growth, and dissipation within a few seconds^30^. It is therefore possible that the membrane receptors at the actin foci site experience transitions in micro- and nano-scale forces during different phases of the foci lifetime, enabling mechanotransduction. Indeed, recent studies have indicated the role of adhesion receptors such as CD2 in physically supporting TCR signaling^55^, and WASP functionally interacts with CD2 protein, indicating that there may be a connection between WASP and adhesion at the immunological synapse^56^. Our study in its current form does not discriminate between these possibilities and future experiments will be required to dissect them. Similarly, understanding the mechanisms by which WASP-dependent actin foci could promote stiffness-dependent TCR activation at a molecular level will require further work. At least two prevalent models of force-sensitive TCR activation exist; one, that force improves TCR-MHCp bond lifetime directly at the nanoscale^8,57^, and two, the adhesion or protrusive forces at the cellular scale could counterbalance the disruptive forces asserted on TCR-MHCp bond within synaptic interface^18,19^. It is tempting to speculate that the presence of actin foci at the nascent TCR clusters could generate protrusive forces that could in turn provide mechanical advantage against the TCR-MHCp bond disrupting forces. Further work will be required to tease apart the nanoscale level and cellular mechanical connection between MHCp -TCR and WASP -actin foci.

With a few exceptions, a majority of target cells in physiology fall within 2.5KPa -10Kpa of stiffness^14^. Although the local stiffness at the immunological synapse site on target cells is hard to estimate directly, it is likely to be much higher than the surrounding non-synaptic areas on the cells. This is because the synapse is an actively assembled cell-cell interface constituting large molecular assemblies further scaffolded by cytoskeletal structures underneath. Keeping this in mind, we included 50 Kpa as the highest substrate stiffness regimen in our assays to assess the mechanosensory response in CTLs. It is further notable that there is still some mechanosensory response observed in WASP-/- cells in the high substrate stiffness range, indicating that other molecular players are required for a full-blown mechanosensory response. Importantly, integrin adaptor protein Talin, which is known to exhibit mechanosensory response^58^, also showed enhanced recruitment at synapse under higher substrate stiffness conditions. Thus, integrin activation could contribute to additional regulation of mechanosensitive activation in CTLs^59,60^. Further characterization of cytoskeletal pathways and their dependence on WASP vs. integrin-based machinery will reveal the mechanistic underpinnings of CTL mechanosensitivity.

Together, the data presented in this report highlights a novel molecular mechanism in modulating early TCR signaling, CD8+ T cell activation, and cytotoxic function as target cell stiffness changes. This is distinct from a previous study that implied the role of WASP in generating actin protrusions and forces at the level of target killing^61^. We show that WASP is crucial in both sensing as well as generating the forces at the synapse and is crucial for the early stages of T cell activation itself before the cytotoxicity step would be initiated. **(Fig. S5B)** The findings would be important for understanding the immunological basis of high tumor incidences seen in WAS immunodeficiency individuals, where WASP function is compromised, and imply that tumor cells may escape CTL-based immunosurveillance in physiology by lowering their cortical stiffness, as is commonly seen during metastasis^62–65^.

## Acknowledgments

We thank J. Burkhardt and Nathan Roy for sharing LifeAct-GFP mice. We thank the Central Bioimaging facility and Central Animal facility at the Indian Institute of Science and the Keck microscopy facility at the Whitehead Institute. This work was supported in part by the Koch Institute Support (core) Grant P30-CA14051 from the National Cancer Institute; Indian Institute of Science Startup fund, DST SERB-POWER grant, Infosys award, and Syngene CSR grant (Sudha Kumari). DJI is an investigator of the Howard Hughes Medical Institute. Srishti Mandal receives a fellowship from IISC and Sayanti Acharya is supported by a fellowship from Joint CSIR-UGC NET Fellowship India. The authors declare no conflicting financial interests.

## Author contributions

SM, MM, PG, SA, YCP, NL, AA, EL, and SK performed the experiments, and DJI and SK provided intellectual input.

## Methods

### T cell isolation, CTL generation, and cell culture

CD8^+^ T cells were isolated from the lymph nodes of WT or WASP-/- C57BL/6 or pmel-1 mice. Lymph nodes from mice were extracted and processed for immunomagnetic isolation of CD8+ T cells with a mouse Easy Sep CD8^+^ T cell isolation kit, using the manufacturer’s protocol. Isolated CD8+ T cells were cultured in RPMI 1640 (Gibco) supplemented with 10%FBS, 5µM β-mercaptoethanol, and 1mM Sodium pyruvate.

For CD8+ T cell activation experiments including flow cytometric analysis of activation-induced modulation of surface markers, cytokine release, and proliferation, isolated cells were activated using Cell Stimulation Cocktail (eBioscience #00-4970-03) or anti-CD3/CD28 coated plates and assessed for the indicated markers 24h post initial seeding. For flow cytometric analysis of surface markers, cells were gated on FITC-CD3, and surface staining with either PerCPcy5.5-CD69, PE-CD11a/CD18 (LFA1 clone H155-78) or APC-CD62L was examined in live cells and analyzed using BD LSR Fortessa and FlowJo 10.5 software. For detection of activation-induced cytokine release using ELISA, a mouse IL-2 detection kit (Thermo Fisher #50-171-92) or mouse IFNγ detection kit (Thermo Fisher #88-7314-22) were used as per the manufacturers’ instructions. For proliferation assay, the CD8+ T cells were loaded with 2.5µM CellTrace CFSE (Thermo Fisher) in PBS at 37°C for 10 min, washed extensively, and cultured in IL-2 free media for 2 days before analysis of CFSE fluorescence using flow cytometry.

For tumor synapse, as well as cytotoxicity assays, granular secretion assays, mechanical measurements on PA substrates, and adoptive transfer experiments, CTLs derived from CD8+ T cells were used. To generate CTLs, mouse CD8+ T cells were seeded in 24 well plates pre-coated with 0.5 µg/ml anti-CD3 and 5 µg/ml anti-CD28 in complete RPMI 1640 medium (containing 1% MEM non-essential amino acids, 1mM sodium pyruvate, 50mM 2-mercaptoethanol, 1% insulin-transferrin-selenium supplement, and 10% fetal bovine serum). The medium was supplemented with IL-7 at a concentration of 1ng/µl and IL-2 at a concentration of 10µg/ml. After two days, cells were transferred to non-coated plates at a density of 1x10^6^ cells/ml in complete RPMI supplemented with 100ng/ml IL-21. On day 5, the CTLs were harvested and used for experiments. For both synapse and cytotoxicity assays, B16 cells were incubated with 100ng/ml of IFNγ overnight, pulsed with 10µM GP100 peptide (25-33; EGSRNQDWL; Anaspec AS-64752) for 1h, washed and utilized for the experiments.

For control experiments addressing the potential effects of Myca. leaching, a CD4+ T cell-dendritic cell co-culture system was used. CD4+ T cells were isolated from OTII mouse lymphoid organs using an EasySep mouse CD4+ T cell isolation kit (Stem Cell #19852). Dendritic cells were derived from the bone marrow of a wild-type C57BL/6 mouse as described previously^66^. The bone marrow-derived dendritic cells were pulsed with 10µM OTII-peptide (OVA 329-337 peptide; Anaspec # AS-64777) for 2 hours at 37°C, and then treated with DMSO or Myca., washed, and eventually co-cultured with OTII cells. Subsequently, the surface expression of markers CD62L and CD69 and cytokine secretion were determined after 24 hours of culture using flow cytometry and ELISA respectively, as described above for CTLs. OTII cell proliferation was also determined after 60 hours of culture using CFSE dilution assay and flow cytometry as described above.

### Atomic Force Microscopy

MFP-3D AFM setup (Asylum Research, CA) equipped with the sharpened silicon nitride cantilever (SNL-10, 0.06 N/m of spring constant provided on a box by Bruker) was utilized for B16 stiffness measurements. The spring constant of the cantilever was calculated by the thermal fluctuation method in water resulting in 0.0575 N/m with the AFM photodetector sensitivity of 68.29 nm/V. The cells (>15 cells in each sample) were seeded onto a 10mm cover glass, which was placed into a petri dish containing 500 µL of HBSS buffer before the experiments. On average, 100 force curves per cell were recorded by the visual placing of an AFM probe on the cell. Each force curve was recorded at 1Hz for determination of Young’s modulus by the Johnson–Kendall–Roberts model (JKR) model and analyzed using Igor Pro post-processing software. The probe diameter was set to 10nm, and the Poisson ratio of the cell was taken to be 0.35, where the mean value of Young’s modulus was obtained from the lognormal fitting curve. Recorded curves were analyzed individually, where calculated numbers were used to build statistical histograms.

### Traction force measurements

Polyacrylamide (PA) gel substrates were generated following the methodology described in^51^. Briefly, 18mm coverslip bottom dishes were cleaned with 100% Ethyl alcohol (EtOH) and washed with MilliQ. Next, 300µl of a solution of 0.75% 3-(Trimethoxysilyl) propyl methacrylate (Sigma Life Sciences, Cat# M6614-25ML) and 0.3% glacial acetic acid in EtOH was then pipetted onto the 35mm glass-bottomed coverslip dishes and incubated for 6 minutes at room temperature. The coverslip dishes were then washed and baked at 50°C for 30 min. 10% acrylamide solutions containing fluorescent beads (FluoSpheres; Invitrogen, cat# F8807) were made with varying bis-acrylamide concentrations as previously described^51^. To create a subsequent thickness of 25µm in the gel, 6.5 µL of acrylamide solution was pipetted onto the coverslip dishes, and clean 18mm coverslips were applied on top. The gels were polymerized at room temperature for 25 minutes before being submerged in MilliQ H2O. Coverslips were then removed and 300µL of 0.2mg/mL sufosuccinimidyl-6-(4’-azido-2’-nitrophenylamino)-hexanoate (sulfo-SANPAH; Pierce Biotechnology) in H_2_O was pipetted onto the surface of the gel. The gels were subsequently UV-treated and washed with H_2_O. Another round of fresh sulfo-SANPAH was applied and the gels were again exposed to UV and washed. After UV exposure, a protein solution of anti-CD3 (10µg/ml) and ICAM1 (2µg/ml) was added to the gels at 37⁰C overnight. Gels were then washed extensively and stored for 2 days at room temperature to allow free acrylamide to leach out, then washed again with PBS before use in experiments^51^. For traction force measurements, activated CTLs suspended in phenol red-free X-vivo-10 media (Lonza) supplemented with 10% FBS were incubated with the PA gel substrates and imaged using Nikon spinning disc confocal microscope equipped with EMCCD. Images were analyzed and quantified using Fiji Software. We computed the displacement of the fluorescent beads at the apical surface of the hydrogel using custom Matlab (Mathworks) code. The RMS traction generated by the cell was calculated based on the displacement field and the known stiffness of the hydrogel substrate.

### Tumor-CTL conjugates

For conjugates imaging, B16 tumor cells were seeded onto coverslips in the presence of 100ng/ml IFNγ for 12h, subsequently loaded with 10µM GP100 peptide for 1h, washed and incubated with pre-activated CTLs for five minutes, fixed, stained and imaged using confocal microscopy. For cytotoxicity measurement, B16 cells were seeded into a 96-well plate in the presence of 100ng/ml IFNγ, loaded with 10µM GP100 peptide, then washed and incubated with CTLs in phenol red-free RPMI media in the presence of 1µM Sytox-red. Acquisition of Sytox fluorescence was used as a marker of cytotoxicity and was measured using a Flex Station 3 plate reader (Molecular Devices), and plotted using Graph Pad Prism software.

### Flow cytometry, pharmacological treatment, immunostaining, and microscopy

For the flow cytometry experiments to evaluate surface activation markers or adhesion molecules, antibodies were purchased either from Biolegend (CD69, clone H1.2F3; LFA1, clone H155-78; CD62L, clone MEL-14; H-2Db, clone KH95; LAMP1) or eBioscience (LAMP1, clone 1D4B). Cells were stained with live dead stain, washed, and then stained with indicated antibodies on ice, washed, fixed, and subsequently analyzed on a flow cytometer. Flow cytometry data were acquired using BD LSRFortessa and analyzed using FlowJo 10.5 software. For imaging experiments with Myca treatment, B16 tumor cells cultured with IFNγ were loaded with 10µg/ml GP100 antigen, treated with 1µM Myca (Enzo) for 10min, washed extensively, and incubated with CTLs subsequently. For imaging synapses, the conjugates were fixed using PHEM buffer (18.14g PIPES, 6.5g HEPES, 3.8g EGTA, 0.99g MgSO4, pH 7.0 w/ KOH in 1L water) with 3.7% paraformaldehyde, and stained overnight with primary antibodies at 4⁰C (phospho-CD3ζ pTyr-142 from Sigma # SAB4301233; Talin from Santacruz # sc-7534; anti-phospho-Zap-70 (Tyr319)/Syk (Tyr352) from Cell signaling technology # 2717S; Alexa-488 Phalloidin from Thermo Fisher # A12379; anti-phospho Y290 WASP from Abcam # ab59278) followed with secondary antibodies (Jackson Immunoresearch) for 2h at room temperature, washed and imaged using Andor spinning disc confocal microscope. For live imaging of CTL F-actin dynamics, CTLs derived from LifeAct-GFP expressing pmel-1 mice were incubated with peptide-loaded and IFNγ-treated B16 cells cultured in phenol red-free DMEM (Lonza) supplemented with 10% FBS on stage at 37°C, and were imaged using Nikon spinning disc confocal microscope.

### Determination of granular secretion from CTLs

For estimation of TCR-induced granular secretion for CTLs, 20 million WT or WASP-/- CTLs suspended in serum-free HBSS buffer were activated using plates coated with anti-CD3 (10µg/ml) for 30 min at 37°C. Culture supernatant was isolated and concentrated 10-fold using 100KD Amicon centrifugal filtration units (EMD Millipore) and analyzed for protease activity using a fluorometric protease activity assay kit (Abcam # ab112152).

### Adoptive transfer of CTLs

For adoptive transfer experiments 2-3 months old WT or WASP-/- C57BL/6 mice were used. Either B16F10 or ova expressing B16F10 was injected at the subcutaneous site. Mice were subjected to lymphodepletion by total body irradiation (5 Gy) on the 5^th^ day of tumor implantation. On day 7, the mice were injected with CTLs (10 million/animal). For preparing CTLs, T cells were isolated from splenocytes using EasySep Mouse CD8+ T Cell Isolation Kit (StemCell) and were activated using anti-CD3 and anti-CD28 followed with IL-7 (1ng/ul) and IL-2 (10ng/µl) supplementation. After two days, cells were transferred to non-coated plates at a density of 1x10^6^ cells/ml in complete RPMI supplemented with 100ng/mL IL-21. On day 4 cells were harvested, washed, and suspended in PBS and utilized for intravenous injections.

### Image analysis and statistics

The images were analyzed using Fiji software. For the determination of fluorescence intensity within the cell, cell boundaries were created by the “ROI” utility of Fiji. Foci were estimated as described previously.^44^ Briefly, a moving window of 1.6X1.6 µm^2^ was used to create a Gaussian blur image of the original raw F-actin images. The Gaussian blur image was subtracted from the raw image to create the foci image. Data were plotted using GraphPad Prism or MATLAB. For statistical analysis, Mann-Whitney non-parametric two-tailed t-test was performed to compare the populations of cells. For flow cytometry and cytokine ELISA assays, statistical significance was determined using paired two-way t-test. In all graphs, unless otherwise mentioned, the dataset was normalized to the mean values of the ‘Control’ population and then plotted as a ‘normalized’ value. Most graphs highlight the Mean ± SEM in the scatter plot of normalized values unless otherwise mentioned.

## Notes

### Competing Interest Statement

The authors have declared no competing interest.

## References

1. Grakoui, A. et al. The immunological synapse: a molecular machine controlling T cell activation. Science 285, 221–227 (1999).

2. Basu, R. et al. Cytotoxic T Cells Use Mechanical Force to Potentiate Target Cell Killing. Cell 165, 100– 110 (2016).

3. Bashour, K. T. et al. CD28 and CD3 have complementary roles in T-cell traction forces. Proc Natl Acad Sci U S A 111, 2241–2246 (2014).

4. Kim, S. T. et al. The alphabeta T cell receptor is an anisotropic mechanosensor. J Biol Chem 284, 31028–31037 (2009).

5. Hui, K. L., Balagopalan, L., Samelson, L. E. & Upadhyaya, A. Cytoskeletal forces during signaling activation in Jurkat T-cells. Mol Biol Cell 26, 685–695 (2015).

6. Feng, Y., Reinherz, E. L. & Lang, M. J. αβ T Cell Receptor Mechanosensing Forces out Serial Engagement. Trends Immunol 39, 596–609 (2018).

7. Judokusumo, E., Tabdanov, E., Kumari, S., Dustin, M. L. & Kam, L. C. Mechanosensing in T Lymphocyte Activation. Biophysical Journal 102, L5–L7 (2012).

8. Liu, B., Chen, W., Evavold, B. D. & Zhu, C. Accumulation of dynamic catch bonds between TCR and agonist peptide-MHC triggers T cell signaling. Cell 157, 357–368 (2014).

9. Hu, K. H. & Butte, M. J. T cell activation requires force generation. J Cell Biol 213, 535–542 (2016).

10. Li, Y.-C. et al. Cutting Edge: mechanical forces acting on T cells immobilized via the TCR complex can trigger TCR signaling. J Immunol 184, 5959–5963 (2010).

11. Liu, Y. et al. DNA-based nanoparticle tension sensors reveal that T-cell receptors transmit defined pN forces to their antigens for enhanced fidelity. Proceedings of the National Academy of Sciences 113, 5610–5615 (2016).

12. Feng, Y. et al. Mechanosensing drives acuity of αβ T-cell recognition. Proc Natl Acad Sci U S A 114, E8204–E8213 (2017).

13. Husson, J., Chemin, K., Bohineust, A., Hivroz, C. & Henry, N. Force Generation upon T Cell Receptor Engagement. PLOS ONE 6, e19680 (2011).

14. Saitakis, M. et al. Different TCR-induced T lymphocyte responses are potentiated by stiffness with variable sensitivity. eLife 6, e23190 (2017).

15. Liu, Y. et al. Cell Softness Prevents Cytolytic T-cell Killing of Tumor-Repopulating Cells. Cancer Res 81, 476–488 (2021).

16. Choi, H.-K. et al. Catch bond models explain how force amplifies TCR signaling and antigen discrimination. 2022.01.17.476694 Preprint at 10.1101/2022.01.17.476694 (2022).

17. Zhao, X. et al. Tuning T cell receptor sensitivity through catch bond engineering. Science 376, eabl5282 (2022).

18. Pettmann, J. et al. Mechanical forces impair antigen discrimination by reducing differences in T-cell receptor/peptide–MHC off-rates. The EMBO Journal 42, e111841 (2023).

19. Siller-Farfán, J. A. & Dushek, O. Molecular mechanisms of T cell sensitivity to antigen. Immunol Rev 285, 194–205 (2018).

20. Limozin, L. et al. TCR–pMHC kinetics under force in a cell-free system show no intrinsic catch bond, but a minimal encounter duration before binding. Proceedings of the National Academy of Sciences 116, 16943–16948 (2019).

21. Tolar, P. Cytoskeletal control of B cell responses to antigens. Nat Rev Immunol 17, 621–634 (2017).

22. Spillane, K. M. & Tolar, P. Mechanics of antigen extraction in the B cell synapse. Mol Immunol 101, 319–328 (2018).

23. Roper, S. I. et al. B cells extract antigens at Arp2/3-generated actin foci interspersed with linear filaments. eLife 8, e48093 (2019).

24. Shaheen, S. et al. Substrate stiffness governs the initiation of B cell activation by the concerted signaling of PKCβ and focal adhesion kinase. eLife 6, e23060.

25. Wan, Z. et al. B cell activation is regulated by the stiffness properties of the substrate presenting the antigens. J Immunol 190, 4661–4675 (2013).

26. Zeng, Y. et al. Substrate stiffness regulates B-cell activation, proliferation, class switch, and T-cell-independent antibody responses in vivo. European Journal of Immunology 45, 1621–1634 (2015).

27. Roper, S. I. et al. B cells extract antigens at Arp2/3-generated actin foci interspersed with linear filaments. eLife 8, e48093 (2019).

28. Pathni, A. et al. Cytotoxic T Lymphocyte Activation Signals Modulate Cytoskeletal Dynamics and Mechanical Force Generation. Front Immunol 13, 779888 (2022).

29. Schwarz, U. S. & Gardel, M. L. United we stand – integrating the actin cytoskeleton and cell– matrix adhesions in cellular mechanotransduction. J Cell Sci 125, 3051–3060 (2012).

30. Kumari, S. et al. Cytoskeletal tension actively sustains the migratory T-cell synaptic contact. EMBO J 39, e102783 (2020).

31. Babich, A. & Burkhardt, J. K. Coordinate control of cytoskeletal remodeling and calcium mobilization during T-cell activation. Immunological reviews 256, 80–94 (2013).

32. Wan, Z. et al. PI(4,5)P2 determines the threshold of mechanical force-induced B cell activation. J Cell Biol 217, 2565–2582 (2018).

33. Wan, Z. et al. The activation of IgM-or isotype-switched IgG- and IgE-BCR exhibits distinct mechanical force sensitivity and threshold. eLife https://elifesciences.org/articles/06925/figures (2015) doi:10.7554/eLife.06925.

34. Hui, K. L., Balagopalan, L., Samelson, L. E. & Upadhyaya, A. Cytoskeletal forces during signaling activation in Jurkat T-cells. Mol Biol Cell 26, 685–695 (2015).

35. Houmadi, R. et al. The Wiskott-Aldrich Syndrome Protein Contributes to the Assembly of the LFA-1 Nanocluster Belt at the Lytic Synapse. Cell Rep 22, 979–991 (2018).

36. Thrasher, A. J. & Burns, S. O. WASP: a key immunological multitasker. Nat Rev Immunol 10, 182– 192 (2010).

37. Li, Y., Bhanja, A., Upadhyaya, A., Zhao, X. & Song, W. WASp Is Crucial for the Unique Architecture of the Immunological Synapse in Germinal Center B-Cells. Frontiers in Cell and Developmental Biology 9, (2021).

38. Sims, T. N. et al. Opposing Effects of PKCθ and WASp on Symmetry Breaking and Relocation of the Immunological Synapse. Cell 129, 773–785 (2007).

39. De Meester, J., Calvez, R., Valitutti, S. & Dupré, L. The Wiskott-Aldrich syndrome protein regulates CTL cytotoxicity and is required for efficient killing of B cell lymphoma targets. J Leukoc Biol 88, 1031–1040 (2010).

40. Kumari, S., Curado, S., Mayya, V. & Dustin, M. L. T cell antigen receptor activation and actin cytoskeleton remodeling. Biochim Biophys Acta 1838, 546–556 (2014).

41. Cannon, J. L. & Burkhardt, J. K. Differential roles for Wiskott-Aldrich syndrome protein in immune synapse formation and IL-2 production. J Immunol 173, 1658–1662 (2004).

42. Spillane, K. M. & Tolar, P. B cell antigen extraction is regulated by physical properties of antigen-presenting cells. J Cell Biol 216, 217–230 (2017).

43. Zhai, Y. et al. Cloning and characterization of the genes encoding the murine homologues of the human melanoma antigens MART1 and gp100. J Immunother 20, 15–25 (1997).

44. Kumari, S. et al. Actin foci facilitate activation of the phospholipase C-γ in primary T lymphocytes via the WASP pathway. eLife 4, e04953 (2015).

45. Janssen, E. et al. A DOCK8-WIP-WASp complex links T cell receptors to the actin cytoskeleton. Journal of Clinical Investigation 126, 3837–3851 (2016).

46. Palacios, E. H. & Weiss, A. Function of the Src-family kinases, Lck and Fyn, in T-cell development and activation. Oncogene 23, 7990–8000 (2004).

47. Kersh, E. N., Shaw, A. S. & Allen, P. M. Fidelity of T Cell Activation Through Multistep T Cell Receptor ζ Phosphorylation. Science 281, 572–575 (1998).

48. Chakraborty, A. K. & Weiss, A. Insights into the initiation of TCR signaling. Nat Immunol 15, 798–807 (2014).

49. Sawada, Y. et al. Force Sensing by Mechanical Extension of the Src Family Kinase Substrate p130Cas. Cell 127, 1015–1026 (2006).

50. Glazier, R. & Salaita, K. Supported lipid bilayer platforms to probe cell mechanobiology. Biochim Biophys Acta Biomembr 1859, 1465–1482 (2017).

51. Tse, J. R. & Engler, A. J. Preparation of hydrogel substrates with tunable mechanical properties. Curr Protoc Cell Biol Chapter 10, Unit 10.16 (2010).

52. Varma, R., Campi, G., Yokosuka, T., Saito, T. & Dustin, M. L. T cell receptor-proximal signals are sustained in peripheral microclusters and terminated in the central supramolecular activation cluster. Immunity 25, 117–127 (2006).

53. Chatila, T., Silverman, L., Miller, R. & Geha, R. Mechanisms of T cell activation by the calcium ionophore ionomycin. J Immunol 143, 1283–1289 (1989).

54. Bieling, P. & Rottner, K. From WRC to Arp2/3: Collective molecular mechanisms of branched actin network assembly. Current Opinion in Cell Biology 80, 102156 (2023).

55. Jenkins, E. et al. Antigen discrimination by T cells relies on size-constrained microvillar contact. Nat Commun 14, 1611 (2023).

56. Badour, K. et al. The Wiskott-Aldrich syndrome protein acts downstream of CD2 and the CD2AP and PSTPIP1 adaptors to promote formation of the immunological synapse. Immunity 18, 141–54 (2003).

57. Das, D. K. et al. Force-dependent transition in the T-cell receptor β-subunit allosterically regulates peptide discrimination and pMHC bond lifetime. Proceedings of the National Academy of Sciences 112, 1517–1522 (2015).

58. Yao, M. et al. The mechanical response of talin. Nat Commun 7, 11966 (2016).

59. Jankowska, K. I. et al. Integrins Modulate T Cell Receptor Signaling by Constraining Actin Flow at the Immunological Synapse. Frontiers in Immunology 9, (2018).

60. Chen, Y., Ju, L., Rushdi, M., Ge, C. & Zhu, C. Receptor-mediated cell mechanosensing. MBoC 28, 3134–3155 (2017).

61. Tamzalit, F. et al. Interfacial actin protrusions mechanically enhance killing by cytotoxic T cells. Sci. Immunol. 4, eaav5445 (2019).

62. Xu, W. et al. Cell Stiffness Is a Biomarker of the Metastatic Potential of Ovarian Cancer Cells. PLOS ONE 7, e46609 (2012).

63. Guck, J. et al. Optical deformability as an inherent cell marker for testing malignant transformation and metastatic competence. Biophys J 88, 3689–3698 (2005).

64. Swaminathan, V. et al. Mechanical stiffness grades metastatic potential in patient tumor cells and in cancer cell lines. Cancer Res 71, 5075–5080 (2011).

65. Liu, Z. et al. Cancer cells display increased migration and deformability in pace with metastatic progression. The FASEB Journal 34, 9307–9315 (2020).

66. P. Matheu, M., Sen, D., Cahalan, M. D. & Parker, I. Generation of Bone Marrow Derived Murine Dendritic Cells for Use in 2-photon Imaging. J Vis Exp 773 (2008) doi:10.3791/773.

