## Supplementary figures and images for "WASP facilitates tumor mechanosensitivity in T lymphocytes"

### Supplemental Figures

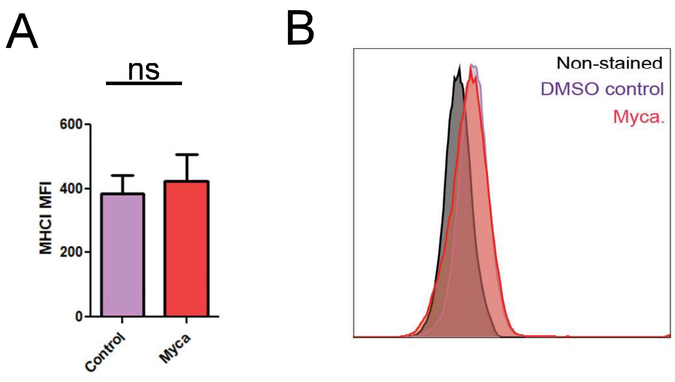

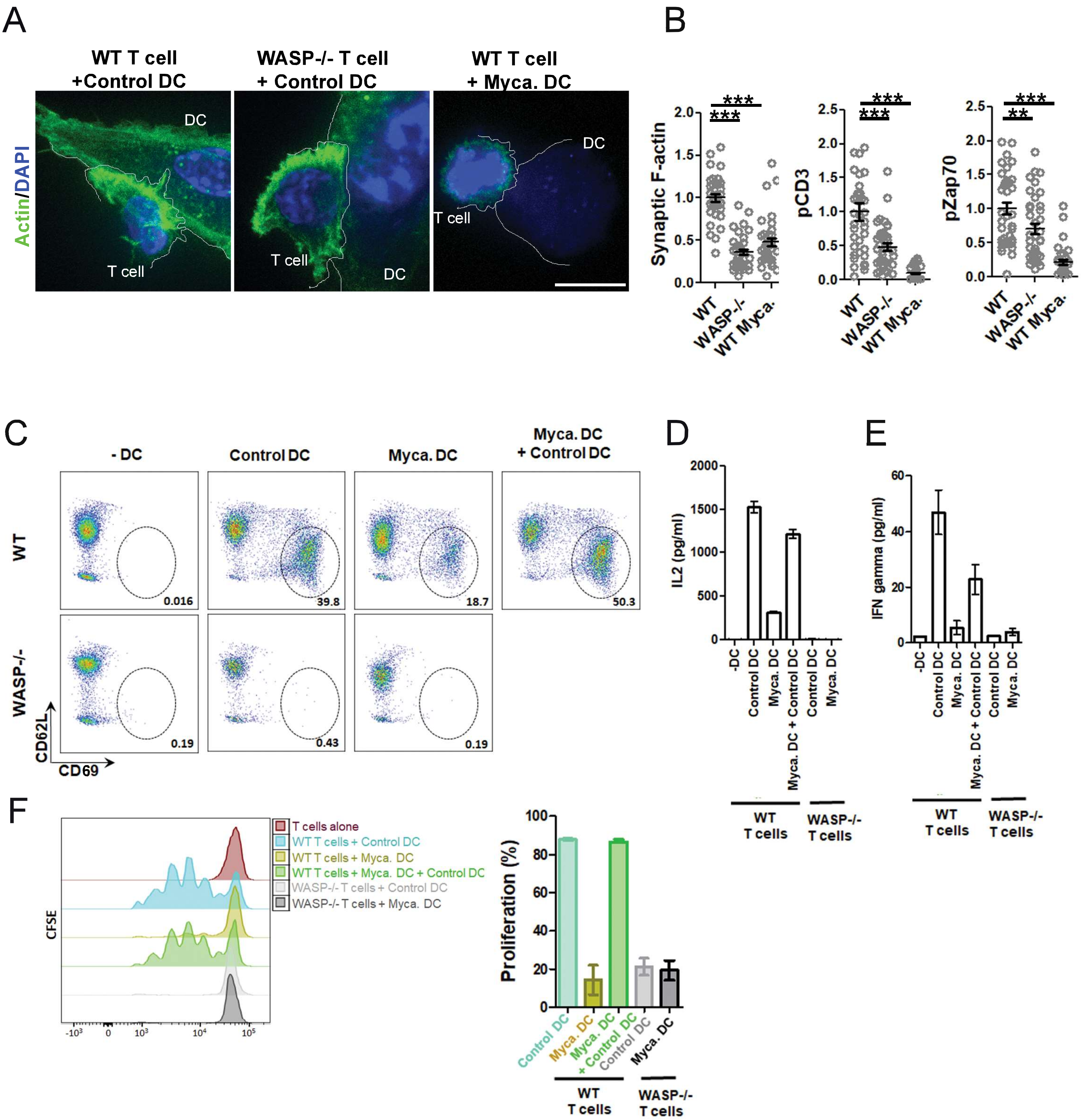

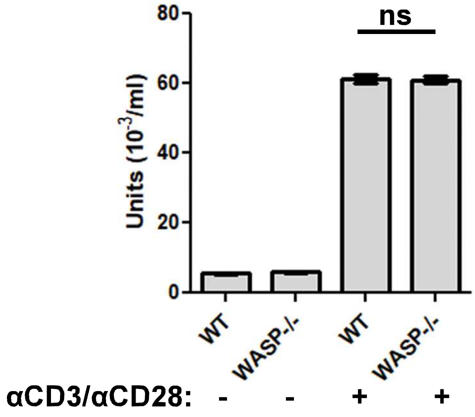

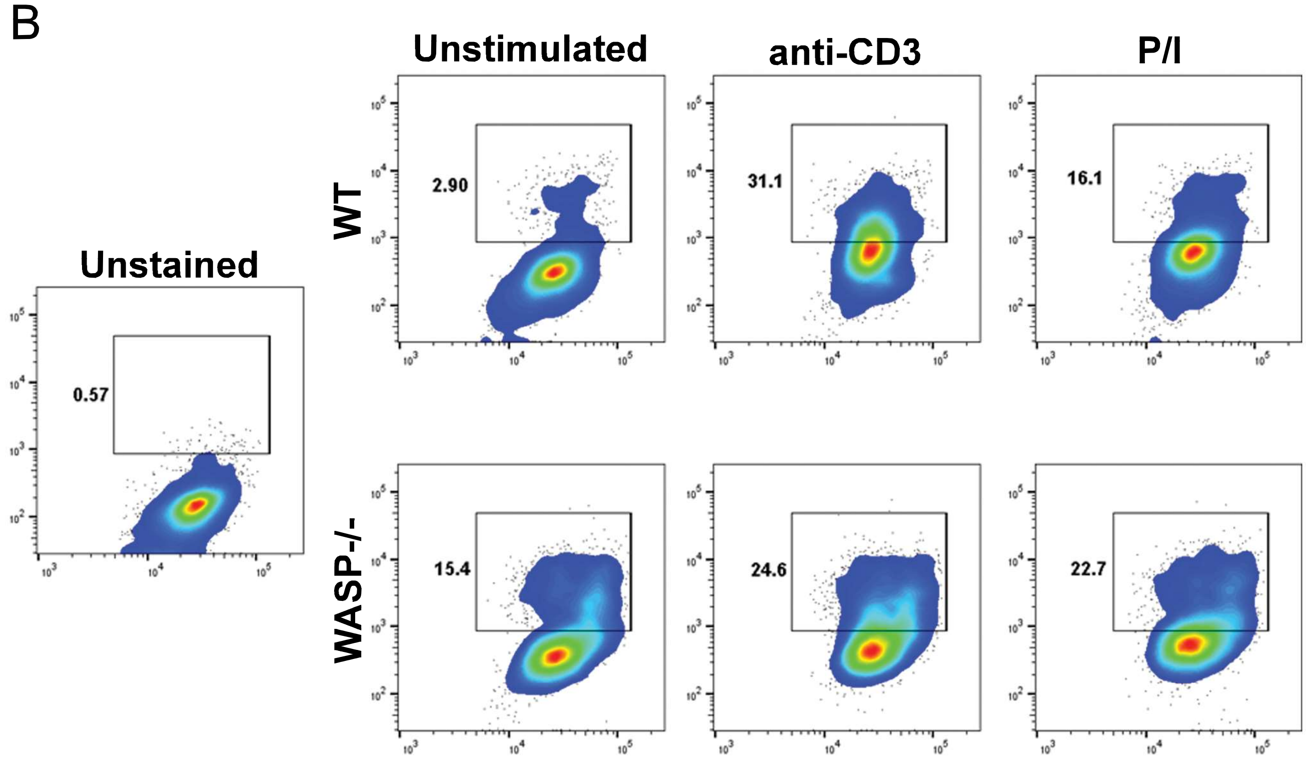

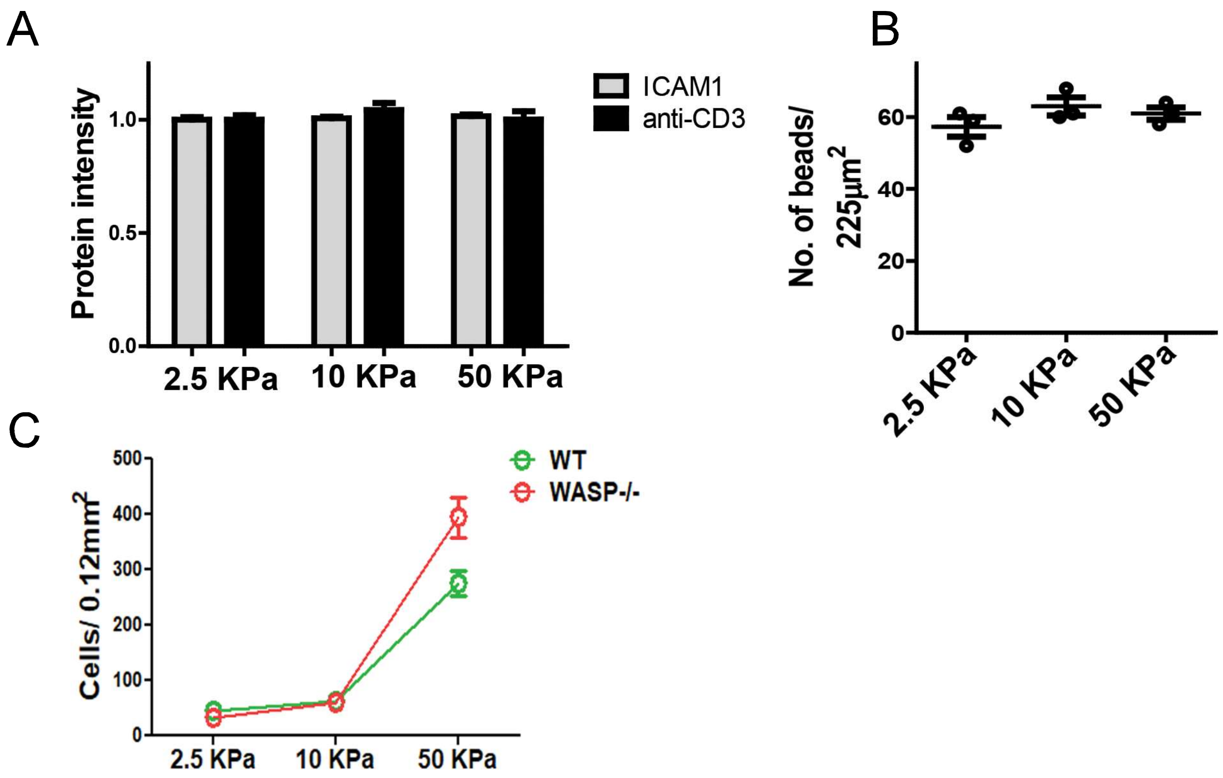

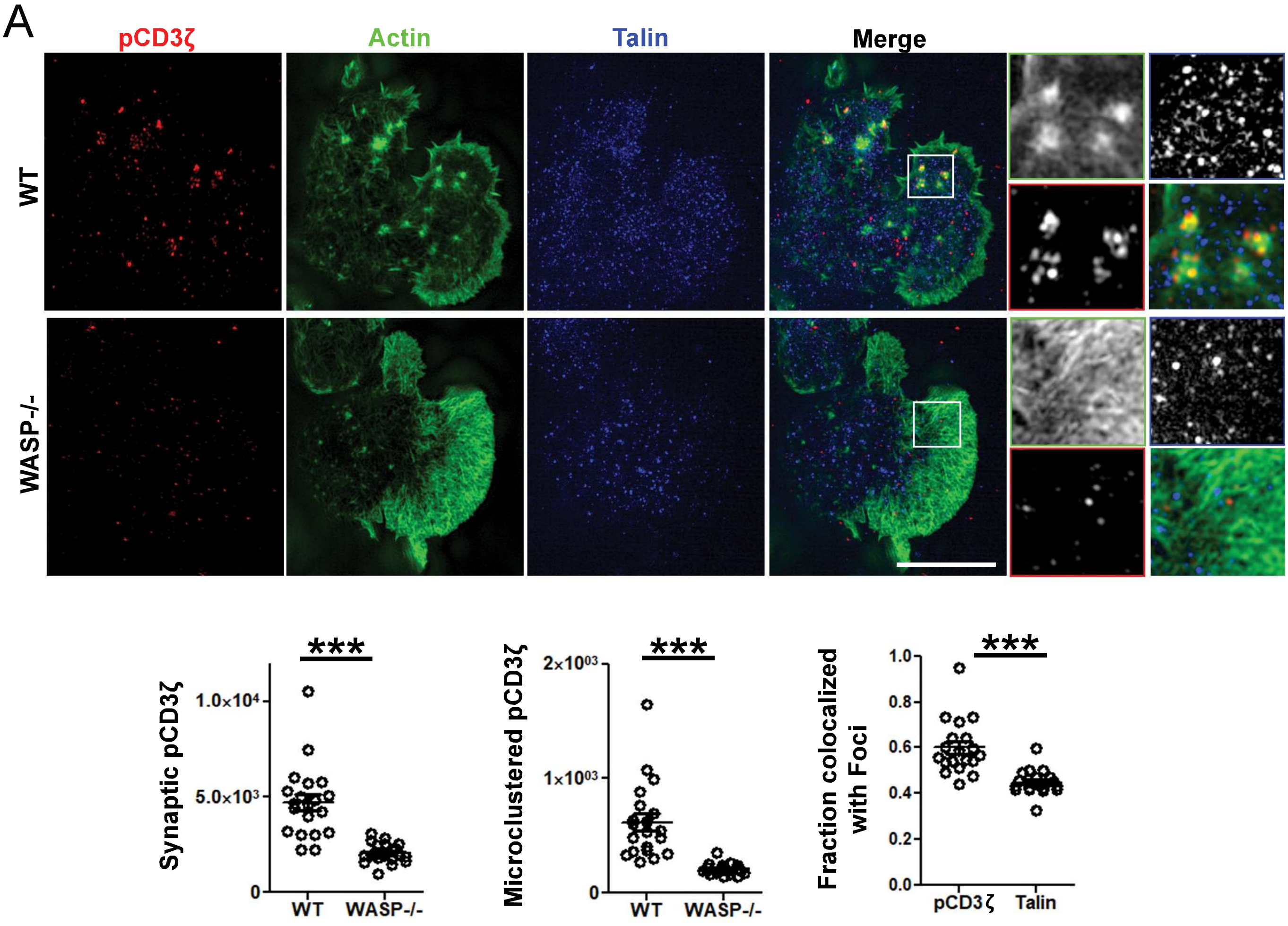

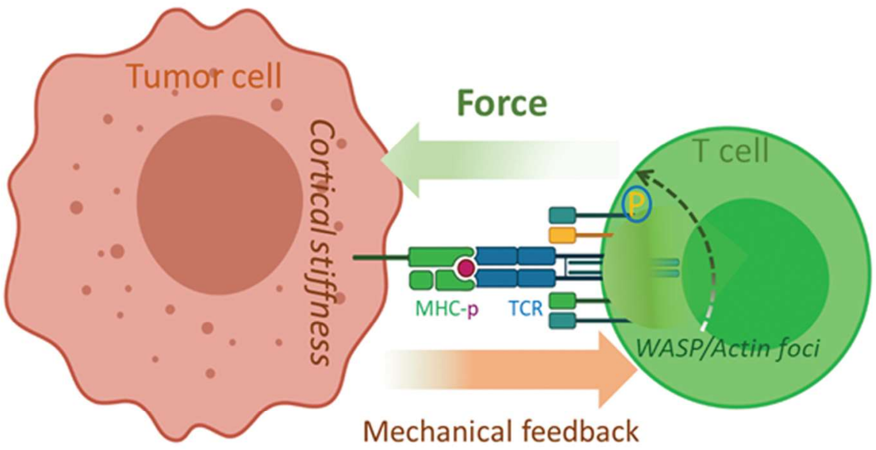
