## Supplemental Text for "WASP facilitates tumor mechanosensitivity in T lymphocytes"

### *"WASP aids early antigen mechanosensitivity in CD8+ lymphocytes"*

#### Supplemental data

**Fig. S1.** (A, B) Surface expression profile of H-2D$^{b}$ on B16 cells assessed using flow cytometry. IFNγ -treated and antigen-loaded B16 cells were further treated with Myca. for 10min, washed and assessed for surface expression of MHCI H-2D$^{b}$. The right plot shows the population distribution of expression; the bar graph on the left shows MFI values obtained from three replicates.

**Fig. S2** Myca. treatment and washout in antigen presenting cells and its effects on CD4+ T cell activation. (A) Myca. treatment of BMDCs impairs synaptic actin distribution in BMDC-OTII cell conjugates. The images are maximum intensity projections of confocal images of the conjugates. (B) Myca. treatment of BMDCs impairs early CD4+ OTII cell activation. Confocal images were analyzed for the amount of F-actin (left graph), pCD3ζ (Middle graph, and p-Zap70 (right graph) in cell-cell conjugate interfaces. Myca.-treated BMDCs incited poor CD69 upregulation and CD62L downregulation on OTII cells (C), poor IL-2 (D) and IFNγ (E ) secretion by the OTII cells, and a poor proliferation of OTII CD4+ T cells (F); all of which could be significantly restored by the presence of DMSO-treated BMDCs ("control DC") in the co-culture system.

**Figure S3.** (A) WT and WASP-deficient CTLs show a comparable TCR stimulation-dependent release of secretory proteases. The bars represent the mean of the values from three replicates ± SEM. (B) Comparison of exocytosis in WT or WASP-/- CTLs. The CTLs were incubated with indicated activation stimuli (either anti-CD3 or PMA/Ionomycin, P/I) for 2h in the presence of anti-CD107- Alexa488 antibody, and cellular CD107 levels were quantified using flow cytometry.

**Figure S4.** (A) PA gels functionalized with Alexa568- anti-CD3 and Alexa 647- labeled ICAM1 were fixed and imaged using a confocal microscope. Fluorescence signal from randomly selected regions was quantified and plotted. (B) Quantification of fluorescent bead incorporation into PA gels shows the equivalent distribution in different stiffness gels. (C) Number of WT and WASP$^{-/-}$ CTLs adhered to the gels after initial seeding and a gentle wash to remove unattached cells.

**Figure S5.** (A) Subcellular cytoskeletal features associated with early TCR signaling. CD3ζ phosphorylation in the synapse of WT or WASP-/- CTLs (images on the top) activated using plate-bound anti-CD3 antibody and ICAM1 and imaged using structured Illumination microscopy. Talin localization was used as a control in the experiment. The graphs show total (left graph), clustered (middle graph), or foci localized (right graph) phosphorylated CD3ζ at the synapse. Phosphorylated CD3ζ clustering was determined as described previously[1]. Each point in the graphs represents values obtained from a single CTL. Scale bar, 5µm.

(B) A model of WASP-dependent mechanosensing and mechanotransduction at the CTL -tumor cell synapse. Higher cortical stiffness of tumor cells triggers better TCR activation, WASP phosphorylation, actin foci, and mechanical forces in the CTL that generates a better CTL activation and tumor cytotoxicity response. However, if WASP is lacking in the CTL or has impaired NPF activity, CTL's rigidity-dependent response of higher early TCR signaling, later activation events, mechanical force generation, as well as cytotoxicity responses, are all compromised. A similar effect is seen when tumor cells are "softer", where the CTLs show lower WASP activation, poor early activation, and tumor cytotoxicity. Thus, WASP and the actin organization it generates at the immunological synapse are crucial for mechanical communication between the tumor cells and the cytotoxic T cells.

**Supplemental Movies:**

**Movie 1.** Live imaging of WT or WASP-/- CTL- B16-F10 tumor interface using spinning disc confocal microscopy. The top panel shows an *en-face* view of LifeAct-GFP distribution in the B16- CTL synapses and the bottom panel shows corresponding bright field images. The numbers in the movie show the time since the beginning of imaging (min: sec).

**Movie 2-3. Time-lapse images of traction forces in the WT and WASP-/- CTL- PA-gel interfaces.** The movies show the distribution of stresses in WT (Movie 2) or WASP-/- (Movie 3) CTL-PA gel interface within 1 min, and 5 min after the initial seeding of cells onto the gels. The play rate of the movies is 30 times faster than the acquisition rate.
